# Mathematics Anxiety Selectively Modulates Attentional and Memory-Related Neural Mechanisms During Numerical Cognition

**DOI:** 10.64898/2025.12.29.696845

**Authors:** Simge Altınok, Sertaç Üstün, Kevser Aktaş, Nihal Apaydın, Metehan Çiçek

## Abstract

Mathematics achievement can be influenced by negative emotions and expectations related to numerical tasks. Individuals with high mathematics anxiety often show poorer numerical performance. This study investigated the neural mechanisms underlying different levels of numerical processing associated with mathematics anxiety using number line estimation and arithmetic verification tasks during fMRI. Participants were classified into high (n=22, age=23.09±2.22) and low (n=24, age=22.67±3.29) mathematics anxiety groups based on detailed screening prior to scanning. Trait and test anxiety were also assessed to capture broader anxiety-related characteristics. Before the fMRI session, participants completed assessments of calculation performance and digit span, and a mock MRI session was used to reduce scanner-related stress. Participants completed task and control conditions for each numerical task during fMRI. Neuroimaging findings were analyzed before and after statistically controlling for trait and test anxiety. Results showed that low mathematics anxiety was associated with greater frontal eye field activity during number-space mapping and stronger supramarginal gyrus activity during arithmetic computation compared with high mathematics anxiety. Controlling for general anxiety revealed a dissociation between mathematics anxiety–specific and general anxiety–related neural effects. Overall, mathematics anxiety selectively influenced attentional and memory mechanisms, whereas broader anxiety processes engaged distinct motor and cognitive systems.

**Key Points:**

- Mathematics anxiety differentially modulates neural activity across distinct numerical processes.
- Mathematics anxiety selectively affects attentional and memory-related mechanisms, independent of general anxiety.
- Neural differences in salience and motor-related regions are better explained by general anxiety rather than mathematics anxiety.

## 1. Introduction

Numerical cognition relies on the ability to represent, compare, and manipulate numerical quantities. This ability begins with a sensitivity to approximate magnitudes that emerges even before language development and is known as the foundational core number system (1). With education, this system becomes integrated with the symbolic number system through the acquisition of number symbols (2). Another critical aspect of numerical cognition concerns the ordinality and relative magnitude of numbers, conceptualized as the mental number line (3,4). This internal spatial representation supports efficient numerosity processing and abstract mathematical reasoning (4–6). The development of mathematical skills depends on the successful coordination of these interconnected numerical processes across multiple representational levels (7).

Neuroimaging studies consistently identify parietal regions as central to numerical representation (8). Supporting this view, number-sensitive neurons have been identified in the parietal cortex (9), and lesion evidence indicates that damage to this region impairs numerical abilities (10). Functional MRI studies further show that deep portions of the intraparietal sulcus (IPS) are selectively recruited during calculation (11). Complementary evidence from single-cell recordings demonstrates that frontal regions also contain number-tuned neurons involved in numerical judgments (12,13). Consistent with these findings, arithmetic tasks reliably engage both frontal and parietal regions (14–16), suggesting that numerical processing relies on a distributed fronto-parietal network (14,17,18). Functional connectivity within this network has been associated with mathematical performance, with higher IPS and fronto-parietal connectivity linked to better achievement in adults (19).

Individual differences in mathematical performance arise from both domain-general and domain-specific factors (20). Mathematics anxiety is a domain-specific affective factor characterized by feelings of fear, tension, and worry associated with numerical activities (21,22). Behaviorally, individuals with high mathematics anxiety show reduced sensitivity to numerical magnitude and poorer arithmetic accuracy or processing efficiency (23–26). Importantly, mathematics anxiety selectively disrupts numerical performance while leaving other cognitively demanding abilities relatively intact, such as reading (25) and semantic processing (27). Converging evidence further indicates that individuals with high mathematics anxiety exhibit heightened physiological arousal in response to complex arithmetic tasks, but not to neutral verbal stimuli (28).

Neuroimaging evidence suggests that the emotional component of mathematics anxiety engages regions involved in affective and threat processing, including the insula (29,30) and amygdala (31–33). Mathematics anxiety has also been linked to altered recruitment of fronto-parietal regions both before mathematical problem solving (34) and during performance (31). Together, these neural findings suggest that mathematics anxiety not only involves affective processing but also has downstream consequences for cognitive resources during mathematical tasks. In line with this view, cognitive accounts propose that anxiety-related thoughts reduce working memory capacity (35,36), an effect that becomes more pronounced as task demands increase (37,38). Reduced processing efficiency in individuals with high mathematics anxiety is particularly evident when tasks require greater inhibitory control (39), and interventions targeting mathematics anxiety have been shown to improve arithmetic performance (40).

Taken together, numerical cognition relies on fronto-parietal and executive control networks, and mathematics anxiety influences cognitive and affective mechanisms that depend on these same systems. However, it remains unclear whether mathematics anxiety affects different levels of numerical processing to the same extent, and whether these neural effects reflect mathematics-specific anxiety or broader anxiety-related traits. Notably, mathematics anxiety shares some characteristics with general anxiety–related factors, including trait and test anxiety (41,42). Previous work has shown that mathematics anxiety can mediate the relationship between trait anxiety and responses to complex numerical tasks, indicating both shared and mathematics-specific effects (28). Consistent with this view, heightened prefrontal activity during mathematical thinking has been reported in individuals with higher anxiety, suggesting that general anxiety may contribute to neural responses observed during numerical tasks (43). Moreover, no fMRI studies to date have directly compared magnitude based tasks, such as number line estimation, with higher-order symbolic arithmetic within the same mathematics anxiety sample, despite their distinct cognitive demands.

The present study examines how mathematics anxiety influences neural activation during number line estimation and arithmetic verification, two tasks that engage distinct levels of numerical processing. Individuals with low and high mathematics anxiety completed both paradigms during fMRI to examine task-specific group differences. Based on prior findings indicating reduced processing efficiency and increased cognitive load in individuals with high mathematics anxiety, we expected group differences to be more pronounced during arithmetic verification. Arithmetic tasks place greater demands on executive control and symbolic manipulation, and may therefore be more susceptible to the disruptive effects of mathematics anxiety than magnitude-based tasks. We further examined whether these neural differences remained after adjusting for trait and test anxiety to clarify the extent to which observed effects reflect mathematics-specific anxiety. We hypothesized that mathematics anxiety would differentially affect neural activation across tasks, particularly within parietal and frontal regions associated with numerical and executive processing.

## 2. Materials & Method

### 2.1. Participants

The study included 46 healthy volunteers aged between 18 and 30 years. Participants were divided into two groups, comprising 22 individuals with high mathematics anxiety (HMA; mean age = 23.09 ± 2.22) and 24 individuals with low mathematics anxiety (LMA; mean age = 22.67 ± 3.29). Mathematics anxiety was assessed using the Math Anxiety Rating Scale-Short Version (MARS-SV) developed by Suinn & Winston (44), with its validity and reliability for the Turkish population established by Baloglu (45).

At the first stage of the study, a mathematics anxiety screening was conducted online. To achieve this, we compiled a set of assessment tools including a demographic form, MARS-SV, a handedness inventory, and trait anxiety and test anxiety inventories into a Google Form. The online survey was announced through university classrooms and student groups, as well as via posters with QR codes displayed across the university campus. A total of 195 volunteers completed the MARS-SV, and the distribution of their mathematics anxiety scores was examined. Participants scoring below the 35th percentile were classified as having low mathematics anxiety, while those above the 65th percentile were classified as having high mathematics anxiety. This extreme-groups approach has been commonly used in previous mathematics anxiety research to maximize group differences and increase sensitivity to anxiety-related effects (26,34,46). All participants were identified as right-handed according to the Turkish adaptation of the Chapman & Chapman (47) Handedness Inventory, established by Nalçacı et al. (48). In addition to demographic characteristics, trait anxiety was assessed in participants using the Spielberger State-Trait Anxiety Inventory—Trait (STAI) (49), while test anxiety was measured using the Spielberger Test Anxiety Inventory (TAI) (50). The STAI is used to evaluate participants’ general anxiety levels, whereas the TAI measures how they feel specifically during written and oral exams.

After the online screening, the study groups were invited to join the fMRI stage. We aimed to recruit participants from non-STEM fields to avoid including individuals who regularly engage with mathematics in their education or daily life. Cognitive assessments were conducted before scanning to evaluate general numerical and working memory abilities. Participants were assessed using the Calculation Performance Test (51), which evaluates mathematics performance across the four basic operations and is structured into five columns, each containing 20 questions. They were asked to complete each column within 30 seconds, following the established protocol for adult groups (52,53). Subsequently, participants were examined using the Digit Span Test from the Weschler Memory Scale-Revised (WMS-R) (54). The Digit Span Test is a verbal measure with two subscales. In the digit span forward task, participants recall numbers in the order presented, whereas in the digit span backward task they recall the numbers in reverse order. Forward digit span primarily reflects verbal short-term memory based on a speech-based representation, whereas backward digit span engages the same verbal store but requires more complex retrieval and manipulation processes, indicating greater involvement of executive control (55).

Prior to the fMRI scanning, participants also completed a 5-minute Mock MRI session and familiarize themselves with the MRI environment to reduce potential stress. During the mock scanning session, we collected participants’ oral self-reports regarding their experience and assessed their readiness for the actual scanning session at the end of the procedure. All participants filled the MRI Safety Checklist before scanning and gave their written permission to join the study. This study was approved by the Clinical Research Ethics Committee of Ankara University Faculty of Medicine on 17 June 2021, with the approval number İ6-406-21.

### 2.2. Task Paradigm

The task paradigm is created using the Psychophysics Toolbox (http://psychtoolbox.org/) on MATLAB (9.7.0.1190202 (R2018a)). Participants underwent fMRI scanning while performing two numerical tasks: a number line estimation task and an arithmetic verification task. In the number line estimation task, a number line ranging from 0 to 200 was presented on a grey background, with a black triangle as a cursor above it. We chose the 0–200 range to include three-digit numbers, so that the numerical magnitudes used here would match the magnitudes used in the arithmetic verification task. In each trial, a randomly selected target number was shown, and participants were asked to estimate the position of the given number on the number line. Target numbers were equally likely to fall below or above the midpoint of the line. Participants moved the cursor to the approximate position of the target number using the left or right buttons and then confirmed their estimate with the middle button. The cursor moved smoothly along a continuous horizontal scale; its displacement was not a fixed or discrete unit per button press. In the number line control task, the procedure was identical to the experimental task, except that the presented number was always either 0 or 200 with equal probability, thereby matching perceptual and motor demands while eliminating numerical estimation demands. In the arithmetic verification task, participants were shown an arithmetic calculation consisting of an addition, subtraction, multiplication, or division problem along with its result displayed on the screen. The arithmetic problems used single, two, or three digit operands, and the resulting answers fell within a two to three digit range across all four operations. In half of the trials the displayed result was correct, and in the other half it was incorrect. Participants judged the accuracy of each result. The numerical matching task served as the control condition and required participants to determine whether two presented numbers were equal. The numbers used in this control condition were two or three digit magnitudes, consistent with the numerical range used in the arithmetic verification task. In half of the control trials the numbers were identical, and in the remaining trials they differed. Participants pressed one button if they judged the arithmetic result to be correct or the numbers in the control task to be equal, and another button if they judged the arithmetic result to be incorrect or the numbers to be different (**Figure 1**).

**Figure 1:**
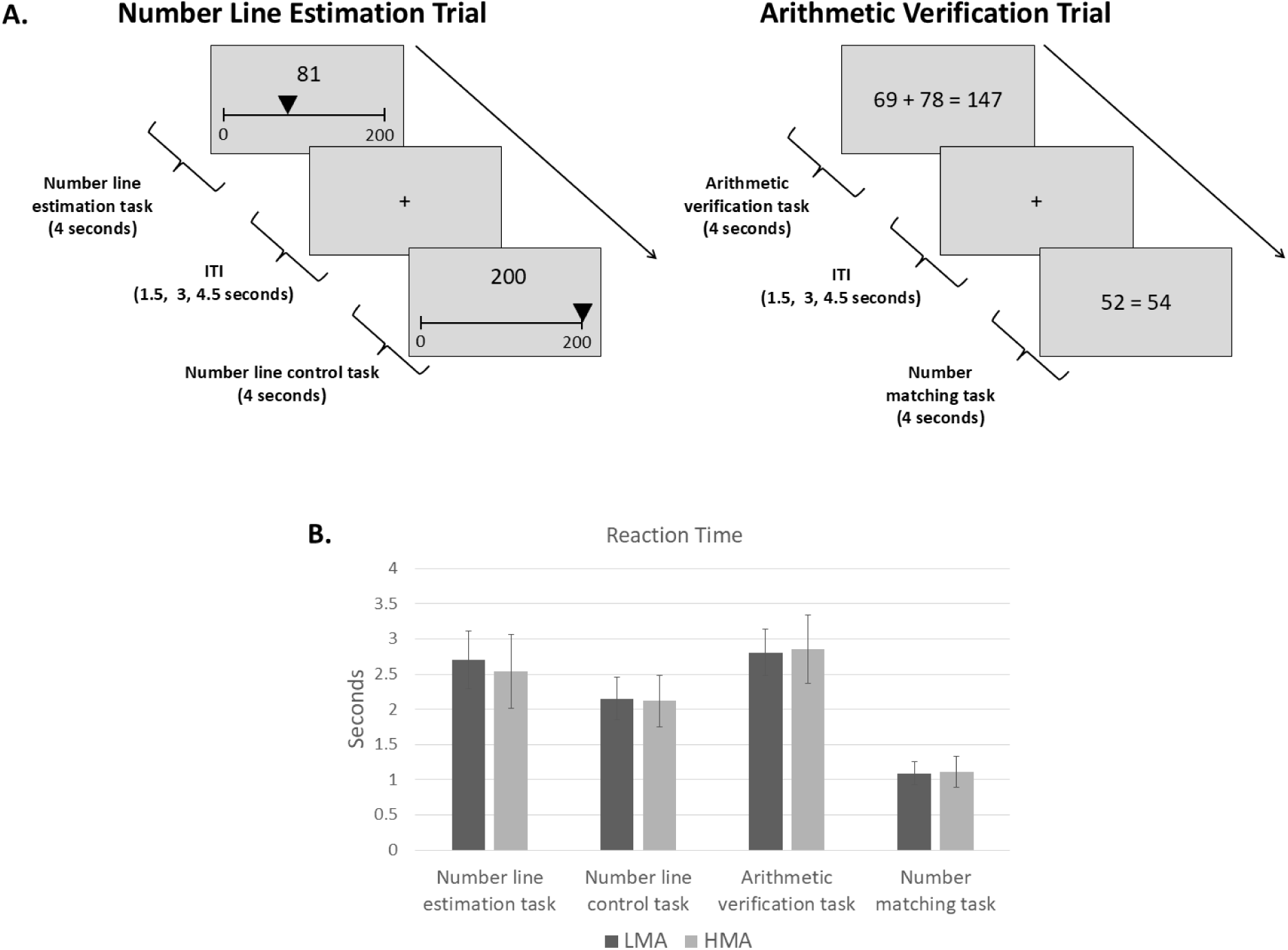
**A.** Experimental task paradigm. **B.** Reaction Times, errors bars represent standard deviations. LMA, low mathematics anxiety; HMA, high mathematics anxiety.

In every task, a fixation cross was presented on the screen between trials for 1.5, 3 and 4.5 seconds in a pseudo-randomized and logarithmic manner that favored shorter durations. An event-related design that consists of four runs was used during scanning. Each run included 40 trials (10 trials per each task condition), with a total duration of 4 minutes and 30 seconds. The four runs consisted of two different types of arithmetic calculations: two involved addition and subtraction, while the other two involved multiplication and division. Two different run orders were employed, starting with either addition–subtraction (A) or multiplication–division (B), resulting in ABBA or BAAB sequences to counterbalance operation type and control for order and fatigue effects across runs. Number line estimation and its control trials were presented in the same event-related runs as the arithmetic and control trials, with all trial types pseudorandomized within each run. Participants were instructed to respond as quickly and accurately as possible. Their responses and reaction times on each task trials were recorded as behavioral data by the stimulus computer via MATLAB which also run the task paradigm during scanning.

### 2.3. Behavioral Data Analysis

In the number line estimation task and its control condition, percentage absolute error (PAE) was calculated. PAE represents the magnitude of the difference between the target value and the estimated value, expressed as a percentage (56). It was computed as: PAE = (|estimate – target| / 200) * 100. To ensure that the number line estimation task reflected numerical magnitude representation rather than response execution alone, we examined whether participants’ estimates were better characterized by linear or logarithmic functions, as commonly assessed in number line estimation research (56). For each model, the resulting R² values (R²_lin and R²_log) were compared using a Wilcoxon signed-rank test. In the arithmetic verification task and its control condition, accuracy was calculated as the percentage of correct responses: Percentage Success = (Number of Correct Answers / Total Number of Trials) * 100. Reaction times were also recorded in all tasks.

All behavioral data collected during scanning were examined using JASP (version 0.18.2.0). Reaction times were analyzed using separate 2 × 2 repeated-measures ANOVAs (group: low vs. high mathematics anxiety; condition: task vs. control) for the number line estimation task and the arithmetic verification task. Similarly, the percentage absolute error in the number line estimation task and percentage of correct responses in the arithmetic verification task were each analyzed using separate 2 (group: low mathematics anxiety vs. high mathematics anxiety) × 2 (condition: task vs. control) repeated-measures ANOVAs. Bonferroni correction was applied to account for multiple comparisons.

### 2.4. fMRI Data Acquisition and Analysis

Before the scanning session began, participants underwent a brief training period to become familiar with the tasks. During scanning, they lay in a supine position inside the scanner. A mirror mounted on the head coil allowed participants to view the task paradigm displayed on an MRI-compatible screen connected to the stimulus computer. A response pad was placed under the participants’ right hand to collect their responses.

A 3 Tesla MRI scanner with a 64-channel head-coil array was used for the image acquisition. High resolution T1-weighted anatomical images had obtained for each participants by the following parameters: Time to Repeat (TR): 2300 ms, Time to Echo (TE): 3.01 ms, Field of View (FOV): 240 mm, slice thickness: 0.9 mm and voxel size: 0.9×0.9×0.9 mm. T2*-weighted functional images were acquired using 48 2.5-mm slices with a 0 mm gap (TR: 1000 ms, TE: 28 ms, Matrix: 84×84, FOV: 208 mm, voxel size: 2.5×2.5×2.5 mm). Each of the four functional runs included 259 TRs, resulting in a total of 1036 images per participant.

Data were preprocessed and analyzed using SPM12 software (https://www.fil.ion.ucl.ac.uk/spm/) run via MATLAB. Functional images were spatially realigned and temporally corrected to the mean of the fMRI images. The functional data were then coregistered with each participant’s anatomical image to enable the localization of activated brain areas. In order to spatially normalize the coregistered data into a standardized anatomical framework, the default EPI template in SPM12 software based on the Montreal Neurological Institute (MNI) average brain and approximating the normalized Talairach coordinates (57) was used. Lastly, functional data were smoothed using Gaussian kernel of 8 mm FWHM.

Stimulus onsets for each trial type (number line estimation, number line control, arithmetic verification, numerical matching) and the six motion parameters were included in the design matrix of the General Linear Model. A high-pass filter of 128 seconds was applied during model estimation. Following the estimation of beta values for the regressors across the four runs, contrast images were generated to represent task-related activation (e.g. number line estimation -number line control and arithmetic verification -numerical matching).

At the second level, the task–control contrast images were entered into separate two-sample t-tests to compare the low and high mathematics anxiety groups for the number line estimation task and the arithmetic verification task. Analyses were conducted both without covariates and with trait anxiety and test anxiety included as covariates, given their conceptual overlap with mathematics anxiety and the significant group differences in these measures. The results were displayed as the directional comparisons between the groups (T contrast; e.g. LMA > HMA and HMA > LMA). Monte Carlo simulation with 10,000 iterations was performed using a voxel-level significance threshold of p < 0.001 to determine a cluster-extent threshold, which was 51 in this study to achieve the correction for multiple comparisons at p < 0.05 (58).

## 3. Results

### 3.1. Demographic and cognitive features of mathematics anxiety groups

Demographic and cognitive characteristics of the mathematics anxiety groups are summarized in **Table 1**. The high and low mathematics anxiety groups were similar in age (*p =* 0.339), gender distribution (*p =* 0.686), years of education (*p =* 0.185), and handedness score (*p =* 0.513). Cognitive test scores, including calculation performance (*p =* 0.062), digit span forward (*p =* 0.982), and digit span backward (*p =* 0.231) were also comparable between the two groups. In line with literature, the high mathematics anxiety group had higher trait anxiety (*p =* 0.021) and test anxiety (*p =* 0.004) scores than the low mathematics anxiety group.

**Table 1.** Sociodemographic and behavioral characteristics of the high mathematics anxiety (HMA) and low mathematics anxiety (LMA) groups.

|  | HMA | LMA | Statistics<br>*p*-value |
| --- | --- | --- | --- |
| All participants (N=46) |  |  |  |
| Subjects (n) | 22 | 24 | - |
| Age (years) |  |  |  |
| Range | 18-28 | 18-30 |  |
| Mean±SD | 23.09±2.22 | 22.67±3.29 | 0.339 |
| Gender (male/female)* | 7/15 | 9/15 | 0.686 |
| Duration of Education (years)** | 16.50±1.83 | 15.67±2.32 | 0.185 |
| Handedness score (right) | 14.00±1.57 | 14.46±2.23 | 0.513 |
| Scales |  |  |  |
| MARS-SV | 88.04±17.18 | 46.83±5.57 | **<.001** |
| STAI-Trait** | 49.36±5.08 | 45.92±4.68 | **0.021** |
| TAI | 42.18±15.75 | 30.33±7.53 | **0.004** |
| Behavioral tests |  |  |  |
| CPT | 95.09±6.05 | 98.33±2.35 | 0.062 |
| WMS-R Digit Span Test |  |  |  |
| Digit span forward | 6.14±1.04 | 6.08±0.83 | 0.982 |
| Digit span backward | 4.91±1.11 | 4.50±1.10 | 0.231 |

MARS-SV: Math Anxiety Rating Scale-Short Version, STAI-Trait: Spielberger State-Trait Anxiety Inventory—Trait, TAI: Spielberger Test Anxiety Inventory, CPT: Calculation Performance Test, WMS-R: Weschler Memory Scale-Revised.
\*Chi-square test was used for nominal data.
\*\*Student's t-test was used for normally distributed data. All other statistical comparisons between groups were performed using Mann-Whitney U Test.

### 3.2. Behavioral findings of fMRI task

#### Number line estimation

For the number line estimation task, percentage absolute error was examined. The ANOVA revealed a significant main effect of condition, *F*(1, 44) = 69.23, *p* < .001, indicating that participants performed worse in the number line estimation task (*M* = 5.94%, *SD* = 2.96) than in the control task (*M* = 0.94%, *SD* = 1.61). Neither the main effect of group, *F*(1, 44) = 1.12, *p* = .337, nor the Group × Condition interaction, *F*(1, 44) = 0.80, *p* = .455, reached significance. Participants’ number line estimates were also better explained by a linear model (*M* R²_linear = 0.918, *SD* = 0.095) than by a logarithmic model (*M* R²_log = 0.834, *SD* = 0.087), as confirmed by a significant difference between model fits (z = 5.851, *p* < .001). A 2 (group: low vs. high mathematics anxiety) × 2 (condition: number line estimation vs. control) repeated measures ANOVA on reaction times revealed a significant main effect of condition, *F*(1, 44) = 109.71, *p* < .001, with slower responses in the number line estimation task (*M* = 2.62 s, *SD* = 0.47) than in the control condition (*M* = 2.14 s, *SD* = 0.33). The main effect of group was not significant, *F*(1, 44) = 0.74, *p* = .395, indicating that overall reaction times did not differ between groups. The Group × Condition interaction was also not significant, *F*(1, 44) = 1.90, *p* = .176, indicating that the size of the task–control difference was comparable across groups.

#### Arithmetic verification

For the arithmetic verification task, a 2 (group: low vs. high mathematics anxiety) × 2 (condition: arithmetic vs. control) repeate measures ANOVA was conducted on both accuracy and reaction time. There was a significant main effect of condition, *F*(1, 44) = 157.54, *p* < .001, with lower accuracy in the arithmetic verification task (*M* = 61.58%, *SD* = 13.64) compared with the numerical matching task (*M* = 98.66%, *SD* = 2.41). Neither the main effect of group, *F*(1, 44) = 2.28, *p* = .115, nor the Group × Condition interaction, *F*(1, 44) = 2.20, *p* = .123, reached significance. Similarly, an ANOVA on reaction times revealed a significant main effect of condition, *F*(1, 44) = 1057.71, *p* < .001, reflecting slower responses during arithmetic verification (*M* = 2.83 s, *SD* = 0.41) compared with the control condition (*M* = 1.10 s, *SD* = 0.19). The main effect of group, *F*(1, 44) = 0.18, *p* = .678, and the Group × Condition interaction, *F*(1, 44) = 0.04, *p* = .843, were not significant. This study used a time-limited arithmetic verification task. Across participants, mean accuracy in the arithmetic verification task was 61.6% and mean reaction time was 2.83 s, indicating that performance was neither at ceiling nor floor and that responses comfortably fit within the 4-second response window.

### 3.3. fMRI findings

#### 3.3.1. Group Differences in Activation During the Number Line Estimation Task

A whole-brain two-sample t-test on the Number Line Estimation > Control contrast revealed greater activation in the left premotor cortex for the low mathematics anxiety group relative to the high mathematics anxiety group **(Table 2**, **Figure 2A)**. When trait and test anxiety were entered as covariates, the pattern of group differences shifted, such that the remaining effect emerged in the right frontal eye field, with the low mathematics anxiety group again showing stronger activation **(Table 3**, **Figure 2B)**. The HMA > LMA comparison yielded no significant activations.

**Figure 2:**
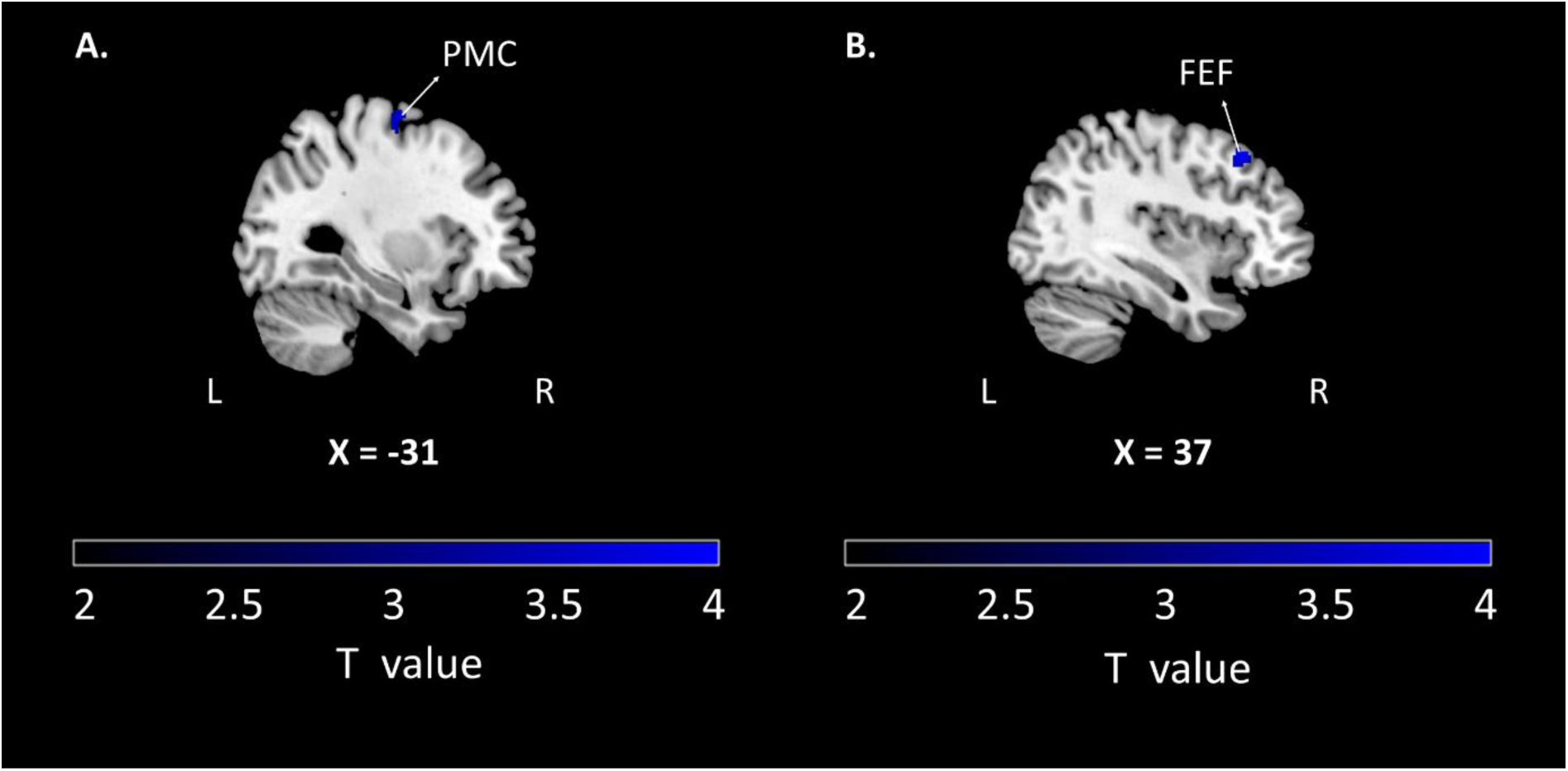
Brain regions showing significant group differences during the number line estimation task **A)** unadjusted model **B)** model adjusted for STAI-Trait and TAI scores (p < 0.001, k ≥ 51, cluster-extent corrected at p < 0.05). PMC: Premotor Cortex, FEF: Frontal Eye Field.

**Table 2.** Brain regions showing significant group differences during the number line estimation task (*unadjusted model*; p < 0.001, k ≥ 51, cluster-extent corrected at p < 0.05).

3.3.1. Group Differences in Activation During the Number Line Estimation Task
| Brain Region | Cluster Size | Laterality | MNI Coordinates |  |  | Z-Score |
| --- | --- | --- | --- | --- | --- | --- |
|  |  |  | X | Y | Z |  |
| ***LMA>HMA*** |  |  |  |  |  |  |
| Premotor cortex | 59 | L | -32 | -10 | 66 | 3.46 |
| ***HMA>LMA*: No clusters survived $k \geq 51$ voxel threshold** |  |  |  |  |  |  |

**Table 3.** Brain regions showing significant group differences during the number line estimation task (*model adjusted for STAI-Trait and TAI scores*; p < 0.001, k ≥ 51, cluster-extent corrected at p < 0.05).

| Brain Region | Cluster Size | Laterality | MNI Coordinates |  |  | Z-Score |
| --- | --- | --- | --- | --- | --- | --- |
|  |  |  | X | Y | Z |  |
| **LMA>HMA** |  |  |  |  |  |  |
| Frontal eye field | 65 | R | 38 | 26 | 46 | 3.65 |
| **HMA>LMA:** No clusters survived $k \geq 51$ voxel threshold | | | | | | |

#### 3.3.2. Group Differences in Activation During the Arithmetic Verification Task

A whole-brain two-sample t-test on the Arithmetic Verification > Numerical Matching contrast revealed greater activation in the left supramarginal gyrus and left lingual gyrus, as well as in the right superior temporal gyrus, right anterior insula, and right anterior cingulate cortex for the low mathematics anxiety group relative to the high mathematics anxiety group **(Table 4**, **Figure 3A)**. When trait and test anxiety were included as covariates, this effect was reduced in extent, with the remaining group difference localized to the left supramarginal gyrus, where the low mathematics anxiety group continued to show stronger activation. No significant clusters were observed for the HMA > LMA contrast **(Table 5**, **Figure 3B)**.

**Figure 3:**
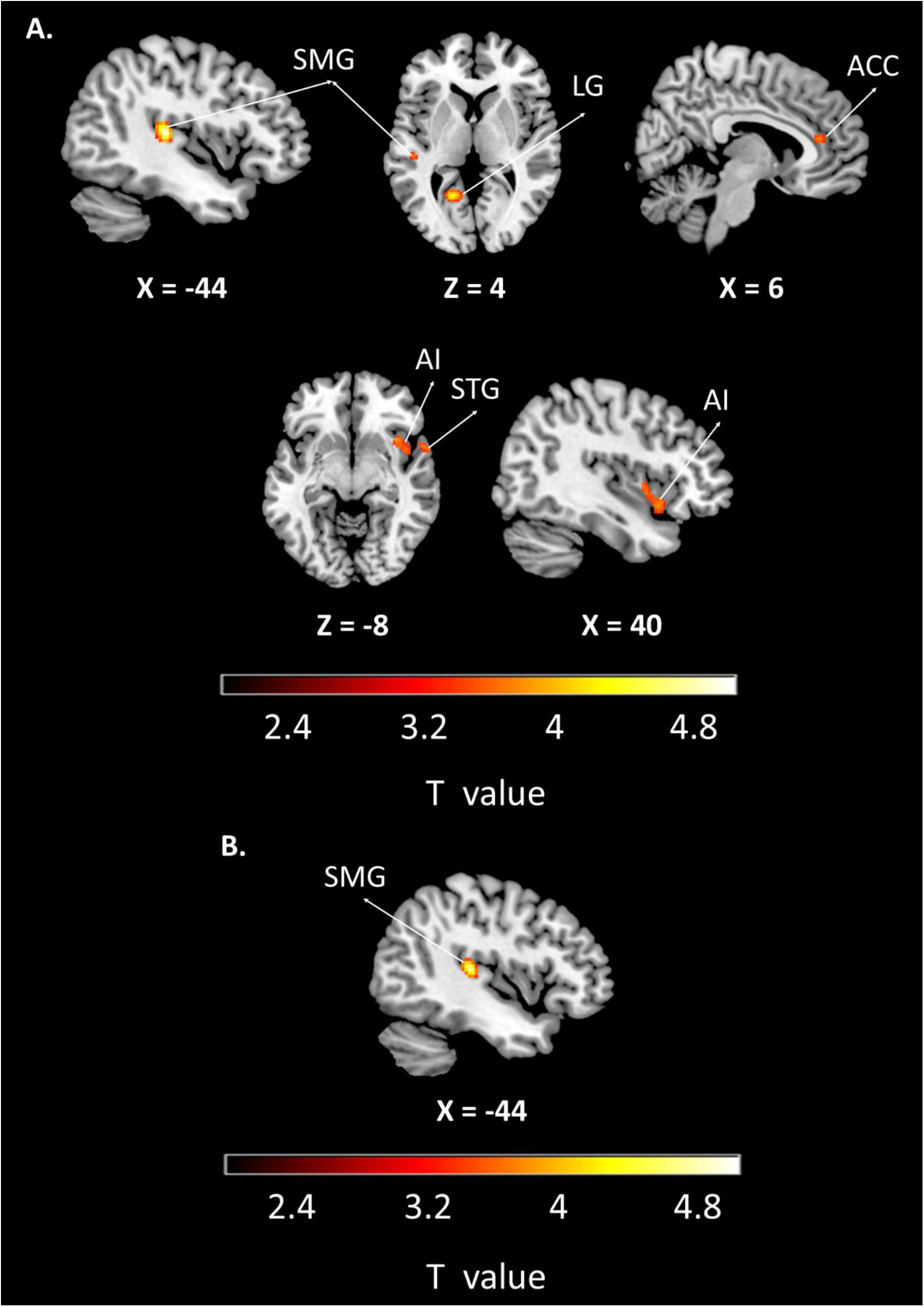
Brain regions showing significant group differences during the arithmetic verification task **A)** unadjusted model **B)** model adjusted for STAI-Trait and TAI scores (p < 0.001, k ≥ 51, cluster-extent corrected at p < 0.05). SMG: Supramarginal Gyrus, LG: Lingual Gyrus, ACC: Anterior Cingulate Cortex, AI: Anterior Insula, STG: Superior Temporal Gyrus.

**Table 4.**
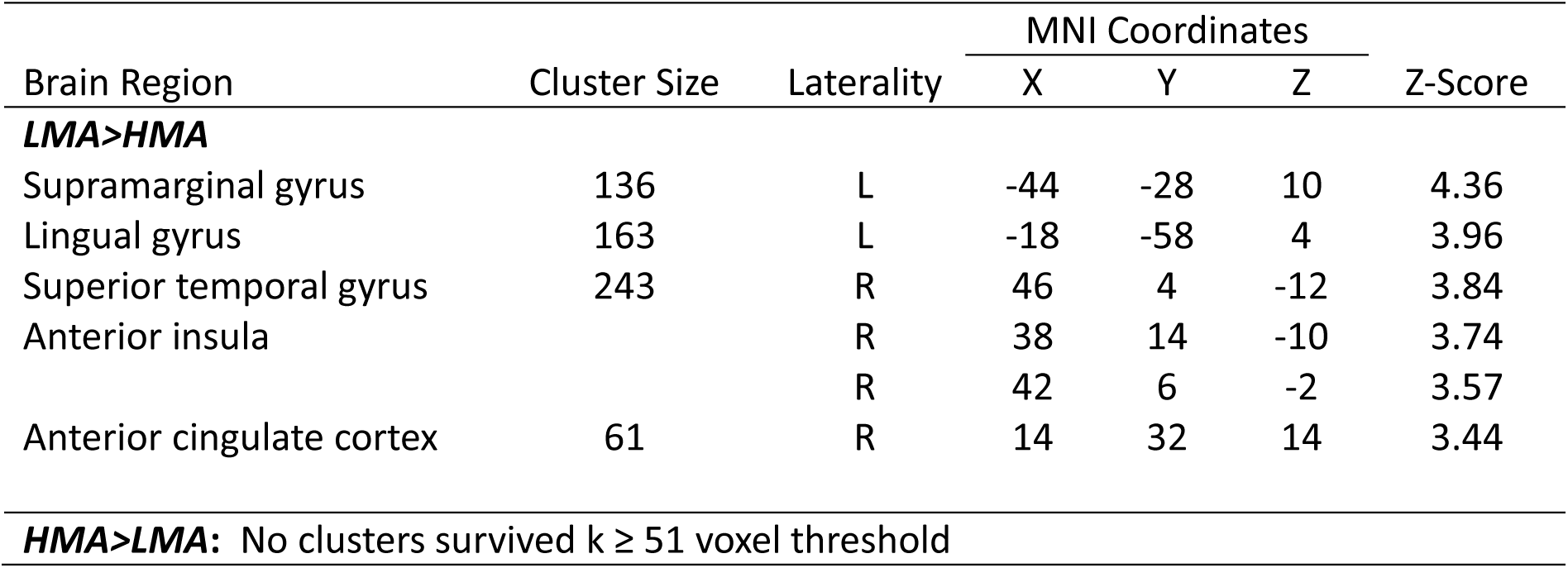
Brain regions showing significant group differences during the arithmetic verification task (*unadjusted model*; p < 0.001, k ≥ 51, cluster-extent corrected at p < 0.05).

**Table 5.**
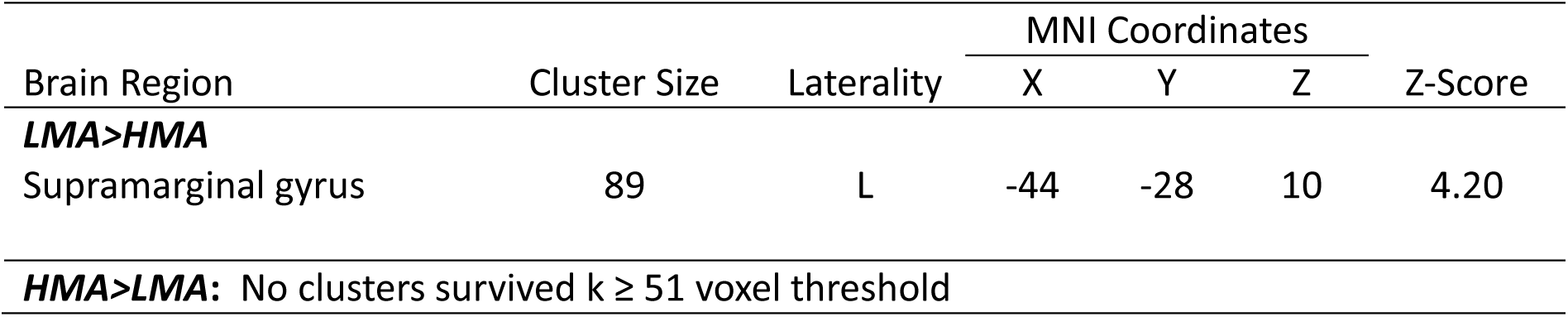
Brain regions showing significant group differences during the arithmetic verification task (*model adjusted for STAI-Trait and TAI scores*; p < 0.001, k ≥ 51, cluster-extent corrected at p < 0.05).

## 4. Discussion

In the present study, we aimed to investigate the neural correlates of different numerical processes in relation to mathematics anxiety. To capture distinct levels of numerical ability, we employed both a number line estimation task and an arithmetic verification task during fMRI scanning. Behaviorally, both the low and high mathematics anxiety groups performed worse in the number line estimation task compared to its control condition, and in the arithmetic verification task compared to the numerical matching condition. Across tasks, participants made more errors and responded more slowly in the experimental conditions. At the neural level, group differences emerged for both tasks and low mathematic anxiety group consistently showed greater activation compared to high mathematic anxiety group in all comparisons which indicate that mathematics anxiety is associated with reduced engagement of task-relevant numerical processing networks. During number line estimation, individuals with low mathematics anxiety showed greater activation in the left premotor cortex compared to those with high mathematics anxiety. After controlling for trait and test anxiety, this group difference shifted, with stronger activation emerging in the right frontal eye field, a region more closely associated with attentional orienting rather than response preparation. Consistent with our hypothesis, the effects of mathematics anxiety were more pronounced during arithmetic verification, a task that places greater demands on executive and symbolic processing. For the arithmetic verification task, individuals with low mathematics anxiety exhibited increased activation in the left supramarginal gyrus, left lingual gyrus, right superior temporal gyrus, right anterior insula, and right anterior cingulate cortex relative to the high mathematics anxiety group. When trait and test anxiety were controlled for, the group difference remained significant only in the left supramarginal gyrus, where the low mathematics anxiety group continued to show greater activation. The persistence of supramarginal gyrus differences after controlling for general anxiety points to a math-specific neural signature associated with arithmetic processing.

### 4.1. Behavioral differences in mathematics anxiety

Even though mathematics anxiety is considered a distinct type of anxiety (42), previous studies have reported correlations ranging from 0.3 to 0.5 between mathematics anxiety and trait or test anxiety, suggesting that these constructs share some common characteristics (21,59). Our results are consistent with this pattern, as we also observed significant differences in trait and test anxiety scores between the mathematics anxiety groups. Given this overlap, we controlled for trait and test anxiety in the fMRI analyses which allows us to dissociate neural effects uniquely associated with mathematics anxiety from those related to more general anxiety processes.

Mathematics anxiety is also known to be associated with lower numerical abilities, including symbolic and non-symbolic number approximation (23,25,26) as well as arithmetic calculation (24,25). In our study, calculation performance showed only a nonsignificant trend (p = 0.062) favoring the low mathematics anxiety group; therefore, behavioral performance was not included as a covariate in the fMRI analyses. Importantly, the absence of robust group differences in task performance during fMRI suggests that the observed neural differences are unlikely to be caused by differences in overall cognitive load. Behavioral task scores during fMRI further revealed that both groups performed more poorly on the number line estimation and arithmetic verification tasks than on their respective control conditions, confirming that the tasks successfully imposed higher numerical processing demands. Another potential cognitive factor linking anxiety and mathematical performance is working memory (35,60). However, in present study the mathematics anxiety groups showed comparable performance on both forward and backward digit span tests, indicating that group differences in neural activation cannot be attributed to baseline differences in working memory capacity

### 4.2. Functional brain differences related to numerical tasks in mathematics anxiety

Dehaene (7) argued that representing and manipulating numerical magnitudes along a mental number line provides a basis for developing arithmetic skills. This numerical–spatial representation captures ordinal relations between numbers and supports efficient magnitude processing and arithmetics (5,6,61). Building on this framework, our neuroimaging findings are interpreted in terms of how mathematics anxiety differentially affects of basic numerical representations and more complex components of numerical processing. Given the established overlap between mathematics anxiety and general anxiety-related factors, we further examine whether observed neural differences persist after accounting for trait and test anxiety to clarify the specifity of the mathematics anxiety-related effects.

In the number line estimation task participants indicated their numerical estimates onto specific spatial locations along a visually presented line. Therefore, it engages on visuomotor coordination within a spatial representation of numerical magnitude. Our results showed that the low mathematics anxiety group exhibited greater activation in the left premotor cortex compared to the high mathematics anxiety group. After controlling for trait and test anxiety, this group difference shifted and and the stronger activation emerged in the right frontal eye field. The premotor cortex has been implicated in action planning during numerical magnitude processing (62), and neurons within this region have been shown to support the spatial representation of goal-directed actions (63). More recent evidence further suggests that the premotor cortex supports the transformation of arithmetic computations into motor responses (64), highlighting its role in linking cognitive operations with action execution. The absence of group differences in premotor area after controlling for broader anxiety measures suggests that this region is more sensitive to general anxiety-related response preparation processes. When variance related to general anxiety are controlled, the core difference related uniquely with mathematics anxiety appear to be more strongly expressed in attentional allocation rather than motor or visuospatial mapping. The frontal eye field is a key region implicated not only in the initiation of eye movements (65) but also in covert attentional orienting and the selection of task-relevant stimuli (66,67). Beyond attentional orienting, the frontal eye field has been shown to support broader cognitive control over visual processing, including attentional allocation and perceptual enhancement (68). These functions are particularly relevant for identifying and marking the correct location on a visually displayed number line, where task performance depends on selectively directing attention to spatial positions and linking them to numerical magnitude. Together, this findings suggest that individuals with low mathematics anxiety engage visual attentional processes to a greater extent during number-space mapping compared to participants with high mathematics anxiety.

The arithmetic verification task engaged the left supramarginal gyrus, left lingual gyrus, right superior temporal gyrus, right anterior insula, and right anterior cingulate cortex more strongly in the low mathematics anxiety group than in the high mathematics anxiety group. After controlling for trait and test anxiety, the left supramarginal gyrus was the only region in which a mathematics anxiety related group difference persisted, with the low mathematics anxiety group continuing to show greater activation. Before accounting for trait and test anxiety, additional group differences were observed in the anterior insula and anterior cingulate cortex, with both regions showing greater activation in the low mathematics anxiety group during arithmetic verification. Together, these regions constitute the salience network, which plays a critical role in detecting salient internal and external stimuli and orienting attention accordingly (69). Through this mechanism, the anterior insula and anterior cingulate cortex are thought to initiate cognitive control–related signaling, particularly during demanding tasks such as arithmetic calculation (70,71). The anterior insula has also been associated with task difficulty and effortful engagement during numerical comparison in several studies (72,73). Both of these regions have been implicated in the pathophysiology of anxiety-related disorders (74). Moreover, activation in the anterior insula has been found to increase with higher trait anxiety scores (75). This pattern of greater insula and anterior cingulate cortex activation in the low mathematics anxiety group may partly reflect general anxiety related influences on task engagement and cognitive control that overlap with mathematics anxiety and are reduced after accounting for trait and test anxiety. In line with this interpretation, previous research suggests that anxiety-related cognitive processes do not necessarily impair performance but may, under certain conditions, enhance motivation and recruit compensatory control mechanisms (76).

Similarly, prior to the controlling for trait and test anxiety, the low mathematics anxiety group additionally showed greater activation in the right superior temporal gyrus than the high mathematics anxiety group during arithmetic verification. Electrophysiological (77) and functional imaging studies (78,79) have reported superior temporal gyrus involvement during mental arithmetic. In addition, cortical surface complexity of the right superior temporal gyrus has been related to numerical intelligence (80). Both increased activity (81) and greater total cortical volume of the superior temporal gyrus (82) have also been associated with generalized anxiety disorder. Taken together, the involvement of the right superior temporal gyrus in the low mathematics anxiety group prior to controlling for trait and test anxiety may reflect wider cognitive engagement during arithmetic verification, that is sensitive to general anxiety levels rather than uniquely to mathematics anxiety.

In contrast, the left supramarginal gyrus showed a robust group difference that remained significant after controlling for trait and test anxiety, highlighting its specific relevance to mathematics anxiety. The supramarginal gyrus has been implicated not only in the manipulation and inhibition of learned verbal sequences (83) but also in the maintenance and updating of working memory representations during numerical problem solving (84). Within the central executive network, it is functionally linked to the dorsolateral prefrontal cortex and is involved in higher-order attentional control (69). Simple arithmetic problems that primarily rely on retrieving solutions from memory rather than effortful computation engage the supramarginal and angular gyri (16,20,85). Lyons & Beilock (29) reported that the negative association between mathematics anxiety and arithmetic performance is evident even before individuals begin performing a mathematics task. Supporting this, it has been reported that individuals with low mathematics anxiety rapidly activate parietal and frontal regions before math trials begin (27). Cognitive strategies used to regulate anxiety have been shown to enhance arithmetic performance in individuals with high mathematics anxiety, an effect accompanied by increased activity in the inferior parietal cortex (40). Furthermore, low mathematics anxiety is found to be associated with greater deactivation of default-mode network regions, indicating higher processing efficiency (39). These findings suggest that individuals with low mathematics anxiety may engage the supramarginal gyrus more strongly during arithmetic tasks, reflecting more efficient memory and attention based processing strategies.

The present study indicates that mathematics anxiety influences neural processing across distinct components of numerical cognition, including numerical–spatial mapping and arithmetic computation. Representing numerical magnitudes on spatial locations was associated with greater right frontal eye field activity, suggesting altered attentional allocation linked to mathematics anxiety, whereas premotor cortex engagement appeared to be driven by general anxiety–related motor or visuospatial processes. Notably, mathematics anxiety–related effects were more pronounced during arithmetic verification, consistent with the greater executive and symbolic demands of this task. During the arithmetic verification task, mathematics anxiety primarily affected activation in the left supramarginal gyrus, which may reflect reliance on mechanisms related to attention and memory, whereas activity in the insula, anterior cingulate cortex, and superior temporal gyrus may reflect modulation of cognitive engagement by broader anxiety-related factors. Together, these findings suggest that mathematics anxiety primarily affects the recruitment of cognitive control resources supporting attention and memory during numerical processing, while additional neural differences observed in salience network regions likely reflect shared variance with general anxiety rather than mathematics-specific mechanisms.

## 5. Limitations

There are several limitations of the present study that should be acknowledged. Although we aimed to standardize participants’ educational backgrounds by recruiting primarily from non-STEM fields, complete homogeneity could not be ensured. As a result, some variability in educational background and everyday numerical experience may have introduced heterogeneity in numerical experiences, which could influence beliefs and attitudes toward arithmetic. Additionally, although gender did not differ significantly between the groups, the number of female participants was nearly twice that of male participants in each group. Given that hormonal fluctuations across the menstrual cycle can modulate stress and anxiety responses, the lack of control for menstrual cycle phase represents a further limitation. Finally, the nature of the arithmetic task itself should be considered. We employed an arithmetic verification paradigm, which requires participants to evaluate the correctness of a presented solution rather than generate an answer themselves. Such tasks are typically associated with lower demands on calculation and working memory processes and may therefore preferentially engage retrieval-based arithmetic strategies. Consequently, this may limit the generalizability of the findings to more effortful forms of arithmetic processing that require higher demands on active computation.

## 6. Conclusion

The present study shows that mathematics anxiety influences neural representations across multiple components of numerical cognition, from numerical-spatial mapping to arithmetic computation. However, these effects were more pronounced when executive and symbolic processing demands were higher, as in the arithmetic task. Our findings reveal that mathematics anxiety selectively modulated attentional and memory-related mechanisms, as reflected in activity within the frontal eye field and supramarginal gyrus, whereas other neural differences involving motor processes and the salience network observed during numerical tasks were better explained by trait and test anxiety. This dissociation underscores the importance of accounting for broader anxiety-related factors when investigating the neural basis of mathematics anxiety.

## Statements and Declarations

### Author Contributions

S. Altınok: conceptualization, methodology, programming experiment, data collection, data analysis, visualization, writing-original draft; S. Üstün: conceptualization, methodology, data analysis, visualization, writing-review and editing; K. Aktaş: conceptualization, methodology, writing-review and editing; N. Apaydın: conceptualization, methodology, writing-review and editing; M. Çiçek: conceptualization, methodology, data analysis, writing-review and editing.

### Data availability statement

The data that support the findings of this study are available on request from the corresponding author. The data are not publicly available due to privacy or ethical restrictions.

### Ethics approval statement

This study was approved by the Clinical Research Ethics Committee of Ankara University Faculty of Medicine on 17 June 2021, with the approval number İ6-406-21.

### Patient consent statement

Written informed consent was obtained from all individual participants included in the study.

## Acknowledgments

We thank the participants for their time and cooperation.

